# Scaling down – Evaluation of start-up and microbial stabilisation dynamics in five parallel laboratory-scale CSTR biogas system

**DOI:** 10.64898/2025.12.23.696146

**Authors:** Lisa Ahrens, Ebba Perman, Anna Schnürer

## Abstract

Laboratory-scale continuous stirred tank reactors (CSTRs) are commonly used to study and optimise anaerobic digestion (AD), yet the time required for such systems to become representative of full-scale processes remains unclear. This study investigated start-up and stabilisation dynamics in five parallel 10-L CSTRs operated as direct scale-downs of a full-scale agricultural biogas plant, using identical inoculum, substrate composition, organic loading rate (2.57 g VS L^−1^ day^−1^), hydraulic retention time (55 days), and temperature (37 °C). The reactors were monitored for 213 days (3.9 HRTs), with one reactor operated for an additional 330 days. All reactors showed highly reproducible behaviour but experienced a transient disturbance, characterised by volatile fatty acid (VFA) accumulation and fluctuating methane production after approximately one HRT. This disturbance was likely caused by a higher effective daily organic loading rate compared to full scale, resulting from once-daily feeding excluding weekends. Increasing the feeding frequency led to VFA degradation and recovery, and stable process performance was achieved after approximately three HRTs, with methane yields of 285 ± 21 NL CH_4_ kg^−1^ VS. The microbial community showed pronounced dynamics during start-up, including major genus-level shifts within *Cloacimonadia* and a transition in aceticlastic methanogens from *Methanothrix* to *Methanosarcina* compared to the full-scale system. Although microbial composition stabilised after about three HRTs, it remained dynamic during prolonged operation and diverged from the original inoculum. These results demonstrate that even direct scale-down reactors require extended stabilisation and may develop microbial communities that differ from the full-scale process, raising questions about their representativeness.

## Introduction

Anaerobic digestion (AD) is an attractive route for production of renewable energy (biogas) and nutrient rich digestates from different type or organic waste streams, such as food waste, wastewater sewage sludge, agricultural residues and animal manure are degraded and converted to biogas (Scarlat et al. 2018; Song et al. 2023). Biogas can be used for production of electricity, heat or vehicle fuel, and the digestate is an excellent organic fertiliser (Lamolinara et al. 2022; Sher et al. 2024). Wet digestion is the most establish commercial AD technology and is typically operated with a total solids (TS) content < 15 % (André et al. 2018; Kougias and Angelidaki 2018), while process with TS content exceeding 15% instead is operated at high-sold digestion conditions (Perman et al. 2024; Shoshaa et al. 2024). Biogas production relies on a set of interlinked biological reactions; hydrolysis, acidogenesis, acetogenesis and methanogenesis, mediated by different anaerobic microorganisms (Santinello et al. 2024; Song et al. 2023). Balances between these microbial steps are essential for good process functioning (Harirchi et al. 2022).The efficiency and stability of the biological process depends on an array of different parameters, including substrate characteristics and operational conditions such as temperature, organic load, hydraulic retention time, and reactor technology (Nkuna et al. 2022; Meegoda et al. 2018; Rabii et al. 2019; Song et al. 2023; Westerholm et al. 2019).

The AD technology is well established in commercial scale operation for conversion of an array of different organic waste materials, but even so many processes are still operating under non optimal conditions and occasionally suffers from process instability and disruptions (Angelidaki et al. 2005; Lienen et al. 2013; Wu et al. 2021). For efficient and stable operation, questions remain to be understood and resolved regarding optimal operation management and links between process operation and microbial function and taxonomy (Santinello et al. 2024; Sarker et al. 2019). To avoid disturbance in full scale, laboratory-scale experiments are frequently used to study and optimise AD processes. These experiments allow for cost-efficient, controlled evaluation of different operational conditions, including both chemical and microbial components (Lüdtke et al. 2017; Schnürer et al. 2016). Numerous lab-scale studies have investigated the effect of different substrates, co-digestion strategies, and operational conditions, such as organic loading rates, hydraulic retention times and temperatures, as well as associated microbial responses (Ahlberg-Eliasson et al. 2021; Bouallagui et al. 2010; Gallert et al. 2002; Liu et al., 2018; Mammo et al 2020; Westerholm et al., 2018). Several factors can influence experimental outcomes of such experiments and must be carefully considered, particularly as they might need to be changed as compared to full-scale conditions. These include the need for substrate pretreatment, change of stirring and feeding frequency and inoculum selection during start-up (Bong et al. 2018; Cazaudehore et al. 2026; Li et al. 2021; Li et al. 2022; Liu et al. 2018; Perman et al. 2024; Song et al. 2023; Schnürer et al. 2016). The number of replicate reactors is also a critical parameter when quantifying differences between treatment and control reactors (Hagen et al. 2014; Lüdtke et al. 2017; Schnürer et al. 2016).

General protocols are available for start-up, operation and evaluation of lab-scale AD experiments (Schnürer et al. 2016). These protocols primarily address operational management and chemical performance of the process, while microbial dynamics are less considered. It is commonly assumed that, following start-up, reactors should be operated for at least three hydraulic retention times (HRTs) after reaching target operational conditions to achieve steady-state reactor performance and a stable microbial community (Pavlostathis & Giraldo-Gomez 1991; Schnürer et al. 2016). During operation of continuously stirred tank reactors (CSTRs) there is always a probability of material spending less time in the reactor than the set HRT (Perman et al. 2022), why multiple HRTs are needed to reach chemical and biological steady state. According to first order dilution kinetics, less than 5% of the original material remains after 3 HRT (Pavlostathis & Giraldo-Gomez 1991), but some studies proposed that even longer time (∼4 HRT) might be needed to reach similar process performance and microbial community structure, depending on the initial inoculum source (Duan el al. 2021)

It is established that laboratory-scale biogas reactors inoculated from full-scale plants require several HRTs to reach steady state, particularly when exposed to new operating conditions. However, it remains unclear whether the same stabilisation period is necessary when laboratory reactors are operated as direct down-scaling of existing full-scale processes, using identical inoculum, substrates, and conditions. Such scale-down systems are widely used to evaluate proposed full-scale changes, yet the time required for them to become representative of the original process is poorly understood. Although similar process performance has been reported, differences in microbial community composition suggest that stabilisation dynamics may differ between direct scale-down reactors and laboratory systems subjected to altered conditions (Perman et al. 2024; Westerholm et al. 2019).

The aim of this study was to investigate the scale down of a full-scale AD process and to determine the time required for stabilisation in laboratory-scale reactors. The reactors were in a next step to be used for evaluation of a new management condition. Five parallel reactors were operated for 213 days (3.9 HRTs), after which one reactor was monitored for additional 330 days. The reactors were inoculated using material from a full-scale biogas plant and operated using the plant’s substrate and overall operational conditions. Process performance, stability, and microbial composition were assessed to evaluate the lab-scale processes.

## Materials and method

### Inoculum and substrates

Inoculum and substrates for the lab-scale experiment were collected from a full-scale agricultural biogas plant, operating with a single CSTR with a total volume of 3600 m^3^ (Lövsta, Uppsala, Sweden). The substrate mix during 2023 consisted of liquid cattle manure (61%), pig manure (29%), wheat flour (7%), silage (2%) and solid manure (1%), based on the wet weight. As for the full-scale plant, the substrate mix was supplemented with an iron chloride solution (20.5-24,3 µl) (Plusjärn S 314, FERALCO NORDIC AB). During the lab-scale experiment the solid substrate was collected twice, while the liquid manure was collected once for the operational period day 1-213, and twice more during the period day 214-543. The liquid manure was kept cold (+4°C) during the experiment, and the solid fraction was frozen (-20 °C) in batches (∼10 kg). One batch at a time was thawed and kept cold (+4°C). The liquid manure was mixed with a kitchen blender before application, to reduce particle size. Chemical composition of the different substrate batches is presented in Table 1.

**Table 1.** Chemical composition of different substrate batches used in the experimental reactors. Values are given as percentage or concentration of wet weight.

|  | **Liquid manure**<br>1<sup>st</sup> batch | **Liquid manure**<br>2<sup>nd</sup> batch | **Liquid manure**<br>3<sup>rd</sup> batch | **Crop residues** |
| --- | --- | --- | --- | --- |
| TS (%) | 5.30 | 5.22 | 4.64 | 68.7 |
| VS (%) | 4.34 | 4.16 | 3.68 | 64.4 |
| Total N (g/L) | 2.7 | 2.9 | 2.6 | 20.6 |
| Organic N (g/L) | 1.2 | 1.4 | 1.1 | 19.9 |
| NH<sub>4</sub><sup>+</sup>-N (g/L) | 1.5 | 1.5 | 1.4 | 0.7 |
| Total C (g/L) | 22.6 | 23.0 | 20.1 | 316.6 |
| Total P (g/L) | 0.47 | 0.44 | 0.32 | 6.54 |
| Total K (g/L) | 1.91 | 2.42 | 1.62 | 11.1 |
| Total Mg (g/L) | 0.39 | 0.44 | 0.39 | 2.87 |
| Total Ca (g/L) | 0.92 | 1.04 | 1.21 | 2.03 |
| Total Na (g/L) | 0.38 | 0.41 | 0.33 | 0.16 |
| Total S (g/L) | 0.27 | 0.34 | 0.25 | 1.55 |

### Experimental setup

Start-up phase: Five 10-L laboratory-scale continuous stirred tank reactors (CSTR) (Belach Bioteknik, Stockholm, Sweden) were used and started with 5 L fresh inoculum each from the full-scale plant. The time between sampling of inoculum and filling the reactors was ca 2 h. The OLR and HRT in all five reactors were set to the same as at the full-scale plant, i.e. 2.57 g VS/L day and 55 days, respectively (Table 2). The systems were operated at mesophilic conditions (37°C). For practical reasons, the reactors were during once per day five days per week during day 1-60. Due to a disturbance in all five CSTRs, as indicated by accumulation of volatile fatty acids (VFA), the feeding was at day 61 changed from five days to six days per week, with the same average organic loading rate. This resulted in decreasing VFA levels and stabilisation in all five reactors. After 213 days four of the five reactors were used for another experiment (data not included in this study), however one reactor continued (control in the new experiment) operating under the same conditions as described earlier. Therefore, this reactor was operated for a total of 543 days.

**Table 2.** Operational parameter of five parallel laboratory-scale processes (B1-D2), mean values with standard deviation for all reactors between day 0 -213 and average data for reactor B1(**) between day 214-543.

|  | **B1** | **C1** | **C2** | **D1** | **D2** | **B1 \*\*** |
| --- | --- | --- | --- | --- | --- | --- |
| OLR<br>(gVS/L day) | 2.57 | 2.57 | 2.57 | 2.57 | 2.57 | 2.57 |
| TS out<br>(%) | 6.49<br>(± 0.01) | 6.53<br>(± 0.01) | 6.71<br>(± 0.01) | 6.51<br>(± 0.01) | 6.59<br>(± 0.01) | 8.31<br>(± 0.01) |
| VS out<br>(%) | 5.15<br>(± 0.01) | 5.16<br>(± 0.01) | 5.32<br>(± 0.01) | 5.12<br>(± 0.01) | 5.23<br>(± 0.01) | 6.49<br>(± 0.02) |
| HRT<br>(days) | 55 | 55 | 55 | 55 | 55 | 55 |
| Weekly average<br>SMP (NL CH<sub>4</sub>/kg<br>VS) | 258<br>(± 57.6) | 257<br>(± 48.2) | 242<br>(± 60.0) | 249<br>(± 36.0) | 243<br>(± 47.2) | 264<br>(± 17.6) |
| Total VFA<br>(g/L) | 1.11<br>(± 1.36) | 1.09<br>(± 1.44) | 1.22<br>(± 2.04) | 0.98<br>(± 1.01) | 1.03<br>(± 1.23) | 0.35<br>(± 0.41) |
| NH<sub>4</sub>-N<br>(g/L) | 2.40<br>(± 0.31) | 2.45<br>(± 0.28) | 2.39<br>(± 0.26) | 2.37<br>(± 0.30) | 2.39<br>(± 0.27) | 2.14<br>(± 0.23) |
| pH | 7.86<br>(± 0.18) | 7.88<br>(± 0.13) | 7.86<br>(± 0.15) | 7.90<br>(± 0.10) | 7.87<br>(± 0.12) | 7.77<br>(± 0.18) |
| Alkalinity<br>(mg CaCO<sub>3</sub>/L) | 13969<br>(± 966) | 13970<br>(± 764) | 13759<br>(± 928) | 14262<br>(± 743) | 13876<br>(± 800) | 11898<br>(± 1613) |
| CH<sub>4</sub><br>(%) | 53.1<br>(± 4.01) | 53.5<br>(± 3.68) | 53.2<br>(± 4.83) | 53.9<br>(± 2.87) | 53.4<br>(± 3.54) | 53.3<br>(± 0.50) |

### Analytical methods

Gas production was measured continuously by built-in gas counters (Belach Bioteknik, Stockholm, Sweden), calibrated using a RITTER Drum-type TG0.5 meter (RITTER Apparateabu GmbH & Co. KG, Bochum, Germany). Gas composition (CH_4_, CO_2_, H_2_S, O_2_, Bal) was analysed using a Biogas 5000 device (Geotech Instruments, Coventry, UK). TS and VS content in substrate fractions and digestate were analysed in triplicate according to standard methods (11 times) (APHA, 1998). Substrate and digestate samples were analysed for content of organic N (SS-ISO 13878), NH_4_^+^-N (Foss Tecator, Application Note, AN 5226, based on ISO 11732) and total C (SS-ISO 10 694) by Agrilab AB (1 time) (Uppsala, Sweden). NH_4_^+^-N measurements in the digestate samples during the startup phase was preformed using LCK 302 Ammonium kit (20 times) (Hach Lange Gmbh, Düsseldorf, Germany), as described previously (Perman et al. 2024). From day 213-543, NH_4_^+^-N measurements in the digestate samples were performed (5 times) according to (Foss Tecator, Application Note, AN 5226, based on ISO 11732). Alkalinity and ratio between VFA and alkalinity (FOS/TAC) were measured weekly on fresh, sieved digested samples as described previously (Perman et al. 2024). The concentration of VFA in the digestate samples was measured by HPLC (Shimadzu 2050 Series, Shimadzu, Kyoto, Japan) as previously described (Tsamadou et al. 2025).

### 16 rRNA gene sequencing

DNA extractions were performed on samples from all five reactors at 20 different time points during the start-up phase (day 1-213), and at 30 additional time points for one of the reactors (B1) during day 214-543. All samples were stored at – 20 ° C until extraction. Extraction was done using the FastDNA Spin Kit for Soil (MP Biomedicals Europe) according to manufacturer’s instructions, with the exceptions that aliquots of 300 µL sample were used and an extra washing step was included to remove humic acids as described in Danielsson et al. (2017) and DNA was eluted using 60 µL DES. DNA concentrations were measured using a Qubit 3.0 Fluorometer. 16S rRNA gene libraries were prepared using primers for amplifications of the V4 region (515’F/806R). The library preparation and sequencing (Illumina Novaseq platform) were performed by Novogene (UK) Company Limited, Cambridge, United Kingdom at three different time points. Processing raw sequences and subsequent data analyses were performed as described previously (Perman et al. 2024).

## Results

### Process performance

Approximately one week after inoculation, the specific methane production (SMP) was between 195-225 NL CH4/kg VS in the reactors. After running the reactor for 213 days the average SMP was 247 (±50) NL CH_4_/kg VS across all reactors (Table 2). After about one HRT (∼55 days), all reactors showed signs of process instability, characterised by increasing VFA concentrations and pronounced fluctuations in the SMP, which ranged between 135-438 NL CH_4_/kg VS between day 50-150 (Fig. 1). Following a change in the feeding regime, from five to six feeding days per week after day 60, while maintaining the same average OLR, VFAs were degraded and process performance stabilised. During days 165–213 (after three complete HRTs), SMP had recovered to stable values of 285±21 NL CH_4_/kg VS (Fig. 1). During this stable phase, the gas had an average composition of 54.1 (±0.5) % CH_4_ and 43.1 (±0.5) % CO_2_, while during days 50-100, the composition fluctuated and the CH_4_ content ranged between 38% and 69% (Table 2, Fig. 1).

**Figure 1.**
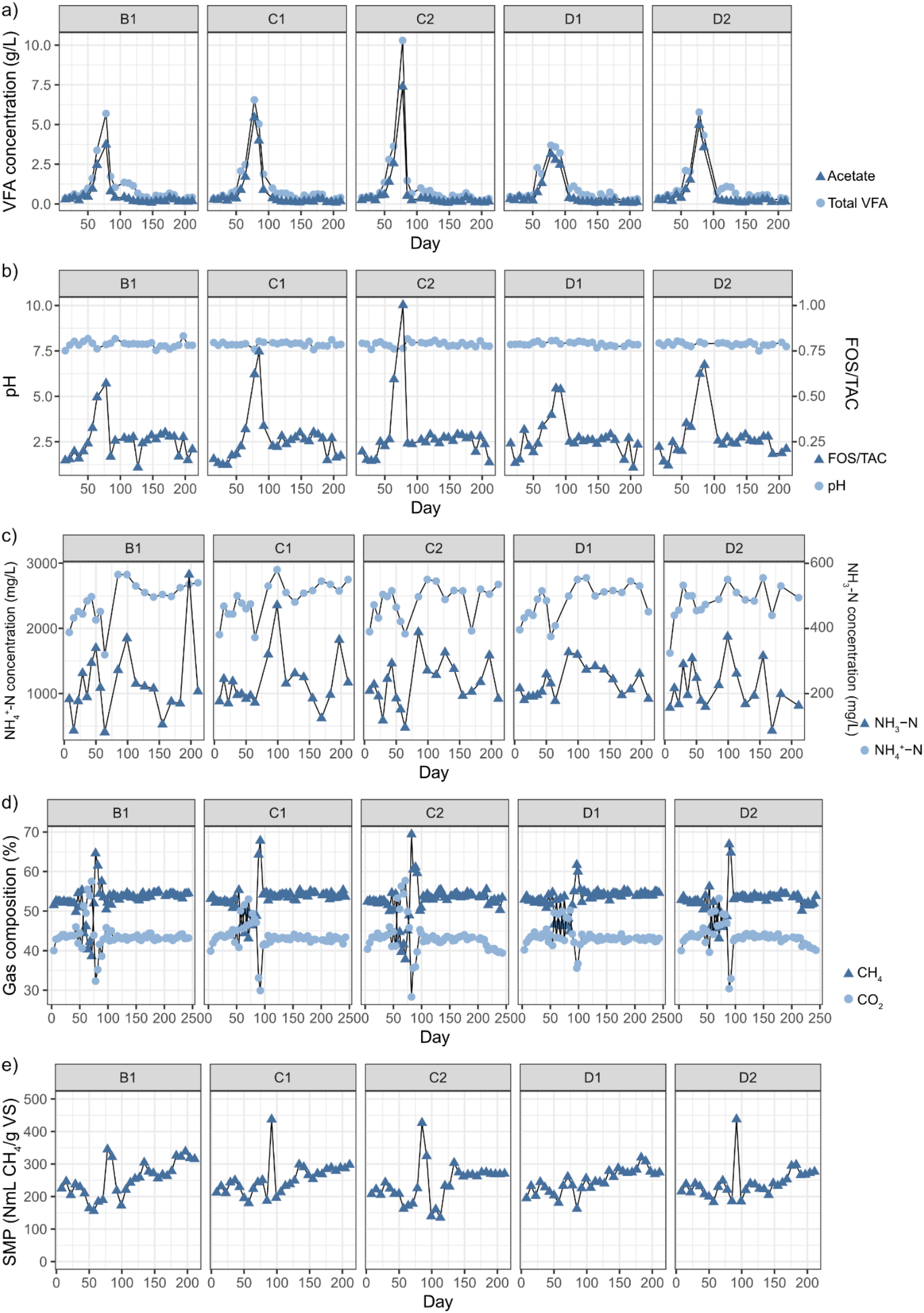
a) Concentration of acetate and total volatile fatty acids (VFA), b) pH and ratio between VFA and total alkalinity (FOS/TAC), c) concentration of NH_4_^+^-N and NH_3_-N, d) percentage of CH_4_ and CO_2_ in gas, and e) specific methane production (SMP), during startup of five parallel continuous stirred-tank reactors (B1, C1, C2, D1 and D2)

The increase in the VFA concentrations, dominated by acetate, became apparent from day 57 and peaked in all reactors on day 78 reaching 5.7, 6.6, 10.3, 3.7 and 5.8 g/L in B1, C1, C2, D1 and D2 respectively (Fig. 1). The VFA concentrations declined rapidly thereafter and remained <1.5 g/L in all reactors from day 100 and onwards. This disturbance was also reflected in the FOS/TAC ratio, which peaked between 0.54-1.00 around day 72-85 (Fig. 1). Outside of this disturbance period, i.e. during the first 50 days after inoculation and between day 100-213, FOS/TAC ratio was generally low (<0.3).

Following the inoculation, the NH_4_^+^-N concentration increased from below 2000 mg/L up to peak with values of 2485-2660 mg/L around day 29-43 (Fig 1). Thereafter, the concentration declined in all reactors, before increasing again around day 85 and subsequently reaching a second peak of 2750-2900 mg/L on day 99 (Fig. 1). From day 100 and onwards, the NH_4_^+^-N levels remained relatively stable generally exceeding 2200 mg/L. Free ammonia-nitrogen (NH_3_-N) followed a similar pattern but showed greater variability during the stable phase, due to pH fluctuations (Fig. 1). Throughout the experiment the NH_3_-N concentrations were within the range 81-565 mg/L.

For reactor B1, operated for an additional period of 330 days, most of the investigated parameters stayed constant, with no significant differences (Table 2). Only TS/VS of the outgoing digestate showed some small differences between the periods.

### Microbial community development

Samples were collected weekly from all reactors for characterisation of the microbial community structure by sequencing of 16S rRNA genes. NMDS-based beta-diversity analysis revealed a gradual shift in microbial community composition throughout the startup phase, which was consistent across all reactors (Fig. 2a). Grouping samples by HRT (1-3) in the NMDS further demonstrated that the community composition changed with each HRT and formed distinct clusters over time (Fig. 2b).

**Figure 2.**
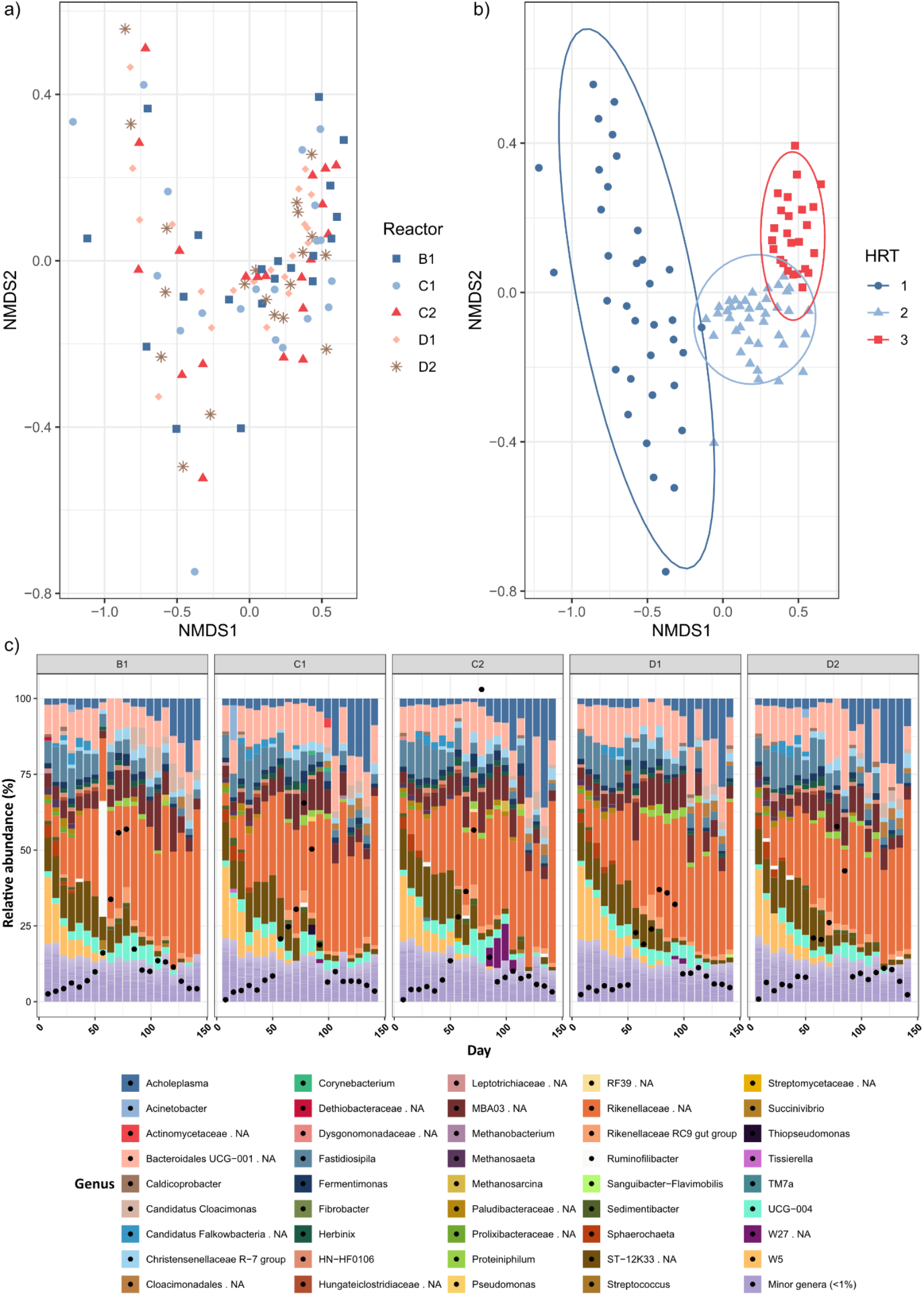
NMDS plots (stress = 0.10) illustrating β-diversity, showing a) reactors B1, C1, C2, D1, and D2, and b) sampling time points grouped by hydraulic retention time (HRT; 55 days) since inoculation. c) Barplot showing relative abundance of taxa on genus level. Black dots indicate concentration of volatile fatty acids (100 on the y-axis corresponds to 10 g VFA/L).

Among Bacteria, the most abundant classes were Bacilli, Bacteroidia, Cloacimonadia, Clostridia and Limnochordia (Fig. S1). Limnochordia was mainly represented by the group MBA03 (RA 3-19%). Within the class Bacilli, two dominant representatives were observed: *Acholeplasma* and UCG-004. *Acholeplasma* had relative abundance (RA) ranging from 1–36%, starting at 2–3% and increasing gradually throughout the startup phase. UCG-004 showed RA values between <1–9%, with its abundance peaking during the VFA accumulation phase after which it decreased to <1% by the end of startup.

Within the class Cloacimonadia, pronounced shifts in genus level composition were observed from inoculation and throughout the startup phase (Fig. S1). At the first time point, genus W5 had the highest RA among the Cloacimonadia as well as the whole community, accounting for 22-24% in all five reactors. However, its RA declined rapidly to 13-18% within one week in all reactors reaching near-zero levels by the end of the startup period. An increase in RA over the startup period was instead observed for a member of the candidate genus *Ca. Cloacimonas*, peaking at RA 4-11%, and an unknown member of the order Cloacimonadales, which reached peak abundance at RA 4-5%.

*Christensenellaceae R-7 group* (class Clostridia) was among the most abundant genera and showed a peak in relative abundance (∼4%) coinciding with the VFA peak, followed by a second increase toward the end of the start-up period across all reactors. In contrast, *Fastidiosipila* (class Clostridia) exhibited high initial RA (7–13%) during the first four weeks, followed by a steady decline over time. The class Bacteroidia was highly abundant in all reactors and increased over time, becoming the dominant group by the end of the start-up phase. This shift was driven primarily by an unclassified member of the family *Rikenellaceae*, which increased from RA 4–6% initially, to 48– 53% at the final sampling point. Another member of the same family, the *Rikenellaceae RC9 gut group*, remained less abundant than its close relative (maximum 4–8%) and the changes in RA followed the VFA dynamics. Also, the Baceroidia ST-12K33 group was initially one of the most abundant genera, present at 11–14% but declined to <3%.

During the start-up phase, the archaeal community shifted from *Methanothrix* (previously *Methanosaeta*) to *Methanosarcina*, coinciding with the peak in acetate (Fig. S1). *Methanothrix* initially comprised 1–3% of the community but declined to near-zero levels after around 8-10 weeks. In contrast, *Methanosarcina* was absent during the first 9–12 weeks but increased after the acetate peak, with RA between 1-3%. A slight increase in *Methanobacterium* was also observed, though its RA stayed below 1%.

### Development of the process and microbiome in stable process

To compare the shifts in the microbial communities during the startup to normal variation within a stable process, one of the reactors (B1) was operated at the same HRT and OLR using the same substrate composition and monitored for a total of 543 days. During this time, samples for microbial community characterisation were taken regularly (Fig. S2). Operational and performance parameters during the extended operation of B1 remained the same (Table 2). However, the microbial community structure showed continued changes. The major changes were observed after change of the batch of liquid manure (day 259 and 371) Specifically, the family *Rikenellaceae* varied in RA, showing an increased RA from around 250 days of operation, with a peak around 300 days, after which the level declined. In contrast, the RA of e.g. Christensenellaceae R7 and Fastidiosipila increased. There was also a slight increase in RA of W5 and *Ca. Cloacimonas* within order Cloacimonadales (Fig. S2). For the archaeal community, an increase in the RA of genera *Methanobacterium* and *Methanosarcina* was seen towards the second half of the experiment (Fig. S2).

## Discussion

Considering chemical characterisation, gas composition, specific methane production (SMP), and microbial community structure, the five parallel reactors investigated in this study exhibited highly similar behaviour. Biogas experiments are generally time- and labour-intensive, and in many cases are therefore conducted using single reactors for both control and experimental conditions. The observed stability and reproducibility among the reactors in the present study suggest that such an approach could be feasible. However, previous studies using parallel laboratory-scale reactors have shown that operational perturbations, such as increases in OLR or ammonia concentration, can elicit divergent responses in microbial community composition across replicate reactors (Lv et al. 2019; Goux et al. 2015). Consequently, the use of parallel reactors remains a robust experimental strategy when responses to a change in operation is evaluated. In the present study, the applied operational conditions may have constituted relatively minor perturbations, as the same substrate and operational parameters were employed as in the full-scale system from which the inoculum was obtained and thus did not trigger diverging responses in the parallel systems.

Members of the order Cloacimonadales decreased in RA as acetate concentration increased, reaching their lowest RA at the VFA peak. Cloacimonadales have been proposed to be involved in propionate oxidation (Alhlert et al. 2016; Pellertier et al. 2008; Westerholm et al. 2021) however, their involvement in acetate metabolism has not been confirmed. Consequently, their specific role in the acetate-associated disturbance observed in the present study remains unclear. Nevertheless, members of Cloacimonadales have previously been identified as indicators of process disturbance inAD systems, as episodes of VFA accumulation frequently coincide with reduced RA of this order (Alhberg-Eliasson et al. 2022; Singh et al. 2021). Possibly the slight increase in ammonia levels as compared to the inoculum could also have been and influencing parameter for the observed shift. Previous studies have shown decreased abundance of Cloacimonadales in response to increasing ammonia levels (Alhberg-Eliasson et al. 2023; Gaspari et al. 2024).

Shortly after inoculation, a high RA of the *Cloacimonadales* group W5 was observed in all reactors. Following the disturbance, however, W5 declined markedly and was replaced by *Ca. Cloacimonas*. One of the major operational changes introduced during downscaling was a modification of the feeding regime, from relatively continuous feeding in the full-scale system to once-daily feeding five days per week at lab scale. This resulted in an increased daily OLR and a shift towards semi-continuous “pulse” feeding, which may have perturbed the microbial community structure. Changes in feeding frequency have previously been shown to induce shifts in microbial population dynamics in AD systems (de Vrieze at al. 2013; Piao et al. 2016). Consistent with this, Perman et al. (2022) reported a decreasing trend in the RA of W5 in reactors operated at higher OLR. Members of *Candidatus Cloacimonas* have also been reported to increase under pulse feeding (Shareki Yekta et al. 2021), which may explain the replacement of W5 by *Ca. Cloacimonas* observed after the disturbance in the present study, as the downscaled feeding regime introduced daily substrate pulses. The observed shift from W5 to *Ca. Cloacimonas* suggests that different *Cloacimonas* lineages may exhibit distinct ecological strategies and tolerance to operational stress, such as elevated OLR and transient VFA accumulation. Such intra-order shifts may therefore reflect functional redundancy within Cloacimonadales, enabling the community to reorganise in response to changing process conditions while maintaining overall system functionality.

Interactions between the methanogens and syntrophic bacteria, such as members of Cloacimonadales may have influenced the observed shift in community structure. Calusinska et al. (2025) recently identified *Methanothrix* as a potential syntrophic partner during propionate degradation by a candidate Cloacimonadota species, suggesting that changes in RA of methanogens may be linked to shifts within the community of Cloacimonadales. The altered feeding strategy following downscaling may also have affected the archaeal community, where a decrease of *Methanothrix* was accompanied by an increased RA of *Methanosarcina* in all reactors. The acetoclastic methanogen *Methanothrix* is known to thrive at low concentrations of acetate whereas *Methanosarcina* grows at higher acetate levels and is also capable of hydrogenotrophic methanogenesis (De Vrieze et al. 2012). Similar reduction in *Methanothrix* abundance in response to decreased feeding frequency, leading to temporarily high loads, and concurrent increase of *Methanosarcina* has been observed before (Piao et al. 2018, Conklin et al. 2006). The slight increase in ammonia could have supported this shift as Methanothix, which is known to be comparably more sensitive than Methanosarcina (De Vrieze et al. 2012. Furthermore, the substrate was grinded before feeding to the laboratory-scale reactors, which may have accelerated the hydrolysis and fermentation rates (Coarita Fernandez et al. 2020), leading to faster production and turnover of acetate. Such conditions could further favour *Methanosarcina*. Notably, acetate peaks in all reactors coincided with the lowest abundance of acetoclastic methanogens, indicating that *Methanosarcina* stabilised the processes when the RA increased, in line with previous observations (Lv et al. 2019b).

Over time, the family *Rikenellaceae* came to dominate the microbial community (Fig. 2 and S2). To date, this family comprises two characterized genera, which are known to ferment glucose to propionate, succinate, and low levels of acetate (Graf 2014) (REF). *Rikenellaceae* are commonly associated with animal gut environments and anaerobic digestion (AD) systems (Koo et al. 2019; Camanaro et al. 2016). Consequently, it is likely that *Rikenellaceae* originated from the cattle manure used as substrate. Previous studies have shown that microbial communities in manure-fed AD processes are strongly influenced by the microbial composition of the feedstock (García Álvaro et al., 2024). For example, the peak in RA observed in reactor B1 around day 300 could be linked to the second batch of liquid manure used day 259-371. Other abundant groups within the same class (Bacteroidia) included the uncharacterised families Bacteroidales UCG-001 and ST-12K33, where the RA of ST-12K33 decreased in parallel with declining acetate concentrations, a pattern consistent with observations by Singh et al. 2021b, who reported reduced abundance of this group in acetate-fed reactors.

Downscaling the AD process inevitably alters several operational parameters, including substrate particle size, mixing regime, feeding strategy, substrate variability, and sampling frequency (Perman et al. 2024). These changes likely contributed to the initial acetate accumulation observed during reactor start-up. However, once the microbial community adapted to conditions favoured by the laboratory-scale setup, the processes stabilised, and no further VFA accumulation occurred when operational parameters were kept constant, even over extended operation (>500 days). Performance indicators such as VFA concentrations and gas composition stabilised approximately 100 days after inoculation, which in this case corresponded to 1.8 HRTs. The stabilisation was likely an effect of both an adaptation of the microbial community and a temporary reduction in OLR to avoid putting pressure on the process, followed by a change in feeding regime from five to six days per week which reduced the daily load. Notably, microbial community composition continued to change over time without affecting process performance, suggesting that community structure alone is not a reliable indicator of process stability without supporting process data.

