## Supplementary Figures for "Scaling down – Evaluation of start-up and microbial stabilisation dynamics in five parallel laboratory-scale CSTR biogas system"

### Supplementary material

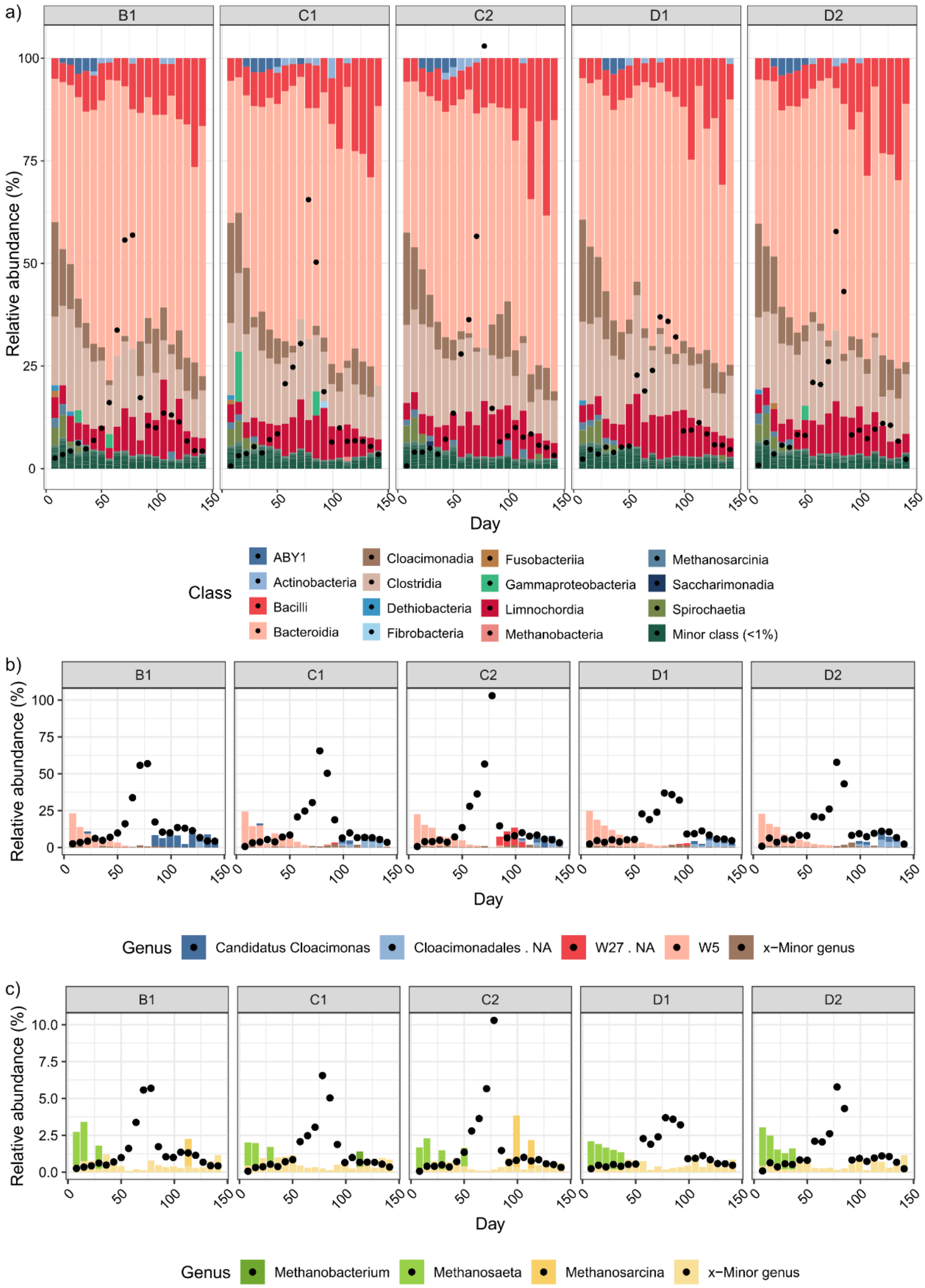


**Figure S1. a)** Barplot showing relative abundance of taxa on class level in reactors B1, C1, C2, D1, and D2. Black dots indicate concentration of volatile fatty acids (100 on the y-axis corresponds to 10 g VFA/L).b) Barplot showing relative abundance of genera within order Cloacimonadales in reactors B1, C1, C2, D1, and D2. Black dots indicate concentration of volatile fatty acids (100 on the y-axis corresponds to 10 g VFA/L). **c)** Barplot showing relative abundance of genera within Archaea in reactors B1, C1, C2, D1, and D2 (plot is zoomed in and shows 0-10% RA). Black dots indicate concentration of volatile fatty acids (10 on the y-axis corresponds to 10 g VFA/L).

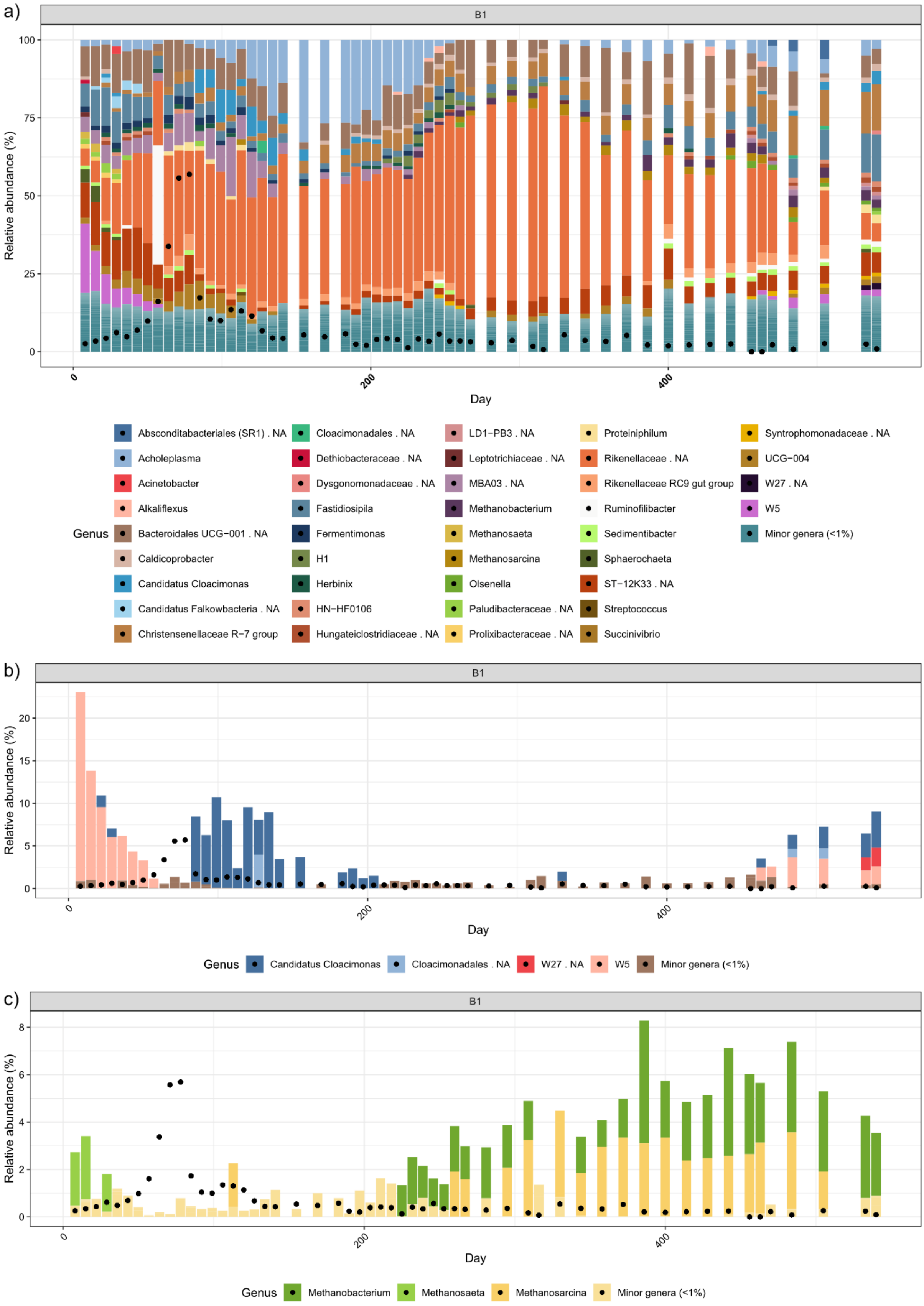


**Figure S2.** a) Barplot showing relative abundance of taxa on genus level in reactor B1 during 540 days of stable operation. Black dots indicate concentration of volatile fatty acids (100 on the y-axis corresponds to 10 g VFA/L). b) Barplot showing relative abundance of genera within order Cloacimonadales in B1. Black dots indicate concentration of volatile fatty acids (10 on the y-axis corresponds to 10 g VFA/L). c) Barplot showing relative abundance of genera within Archaea in B1. Black dots indicate concentration of volatile fatty acids (10 on the y-axis corresponds to 10 g VFA/L).
